# Setting Climate Targets: The Case of Higher Education and Research

**DOI:** 10.1101/2024.03.11.584380

**Authors:** Anne-Laure Ligozat, Christophe Brun, Benjamin Demirdjian, Guillaume Gouget, Emilie Jardé, Arnaud Mialon, Anne-Sophie Mouronval, Laurent Pagani, Laure Vieu

**Affiliations:** Université Paris-Saclay, CNRS, ENSIIE, Laboratoire Interdisciplinaire des Sciences du Numérique, 91400, Orsay, France; Sorbonne Université, CNRS, Institut des Nanosciences de Paris, UMR7588, F-75252 Paris, France; Aix Marseille Univ, CNRS, CINAM, Marseille, France; Université de Rennes, CNRS, ISCR – UMR 6226, F-35000 Rennes, France; Univ Rennes, CNRS, Géosciences Rennes, UMR 6118, 35000, Rennes, France; CESBIO, Université de Toulouse, CNES/CNRS/INRAE/IRD/UT3, 18, Avenue Edouard Belin, 31401 Toulouse, France; Université Paris-Saclay, CentraleSupélec, ENS Paris-Saclay, CNRS, Laboratoire de Mécanique Paris-Saclay, 91190, Gif-sur-Yvette, France; LERMA & UMR8112 du CNRS, Observatoire de Paris, PSL Research University, CNRS, Sorbonne Université, F-75014 Paris, France; IRIT, CNRS, Université de Toulouse, France

## Abstract

The carbon footprint and low-carbon strategies of higher education and research organizations have been the subject of scientific articles and reports. However, these provide few details on the reduction targets themselves, leaving the question of how should higher education and research organizations define and construct their climate targets and trajectories unanswered. The present paper fills this gap. We first review and analyze the documents describing the climate strategies of 53 higher education and research organizations coming from 11 countries, based on their detailed GreenHouse Gas emissions (GHGs) reporting. The selected reports include at least one target reduction for at least one target year. Then, on the basis of this analysis we propose guidelines to encourage and help higher education and research organizations set relevant climate targets.

## 1. Introduction

Like any other institution, universities and other higher education and research organizations must participate in the global effort to reduce GreenHouse Gas emissions (GHGs). The carbon footprint and low-carbon strategies of higher education and research organizations have been the subject of scientific articles and reports [Robinson et al., 2015, Valls-Val and Bovea, 2021, Helmers et al., 2021, ALLEA, 2022]. However, the GHG reduction target itself is rarely discussed: What scopes and emission sources are considered, and how does the target compare to the national ones in terms of reduction percentage and deadline?

This paper discusses the existing targets set for higher education and research organizations, and proposes guidelines to encourage and assist organizations to declare relevant climate objectives. We consider that climate targets benefit from being incorporated into a broader framework of prospective scenario-building, but we will only address the quantitative aspects here.

Thus, our research question is: How should higher education and research organizations define their climate targets and trajectories?

Our contributions are the following:

- Review and discussion about existing targets in higher education and research.
- Guidelines to help higher education and research organizations set climate targets.

Among the existing reports, the European federation ALLEA [ALLEA, 2022] reviews the carbon footprints and reduction strategies of European institutions, including universities and research institutes. We agree with many of the report’s conclusions, in particular, that the absence of a standard on how to report carbon footprints and reduction commitments, makes it difficult to compare institutions and their strategies. In the present paper, we detail the reduction commitments, which are little explored in the ALLEA report.

[Robinson et al., 2015] assess the carbon footprint reduction commitments made by UK higher education institutions and show that the evolution of their carbon footprints does not align with the commitments. They underline that these institutions have not taken the measure of the changes required by the commitments -which were subject to a reporting obligation - and regret the lack of standards concerning scope 3 emissions. [Valls-Val and Bovea, 2021] searched for papers presenting the carbon footprint of higher education institutions and analyzed the data from the 35 papers they identified. Like the ALLEA report, they show that the results are highly diversified and difficult to compare due to the variety of calculation methodologies, temporal perimeters, chosen system boundaries, scopes and items taken into account, and emission factors.

Existing references therefore tend to focus on the carbon footprint of organizations and the measures implemented to reduce this footprint, but few details are given on the reduction targets themselves, which is what we explore in this paper.

Our recommendations in Section 3 are in line with the general guidelines of the the Science Based Targets initiative (SBTi) ‡, which provides organizations “with a clearly-defined path to reduce emissions in line with the Paris Agreement goals”. Many initiatives aim at helping organizations, outside of the higher education and research domain, define their target GHG reductions, the SBTi being a most general one.

## 2. Results

In the following we refer to universities, research laboratories or research institutes with the generic term “organizations”. For the year 2022, we analyzed existing climate strategies of higher education and research organizations from 11 countries and 53 organizations. We relied mainly on public sustainable development policy documents containing a detailed GHG reduction strategy, except for some laboratories participating in the Labos 1point5 collective *§*, for which we had more information at our disposal via the Labos 1point5 experimentation scheme. Figure 1 shows the selected number of organizations per country. More details on our methodology are given in section 5, in particular regarding the way we analyzed and compared the various organizational perimeters in terms of scopes and GHG emission sources, target years, target reductions, and their implementation in GHG reduction trajectories.

**Figure 1.**
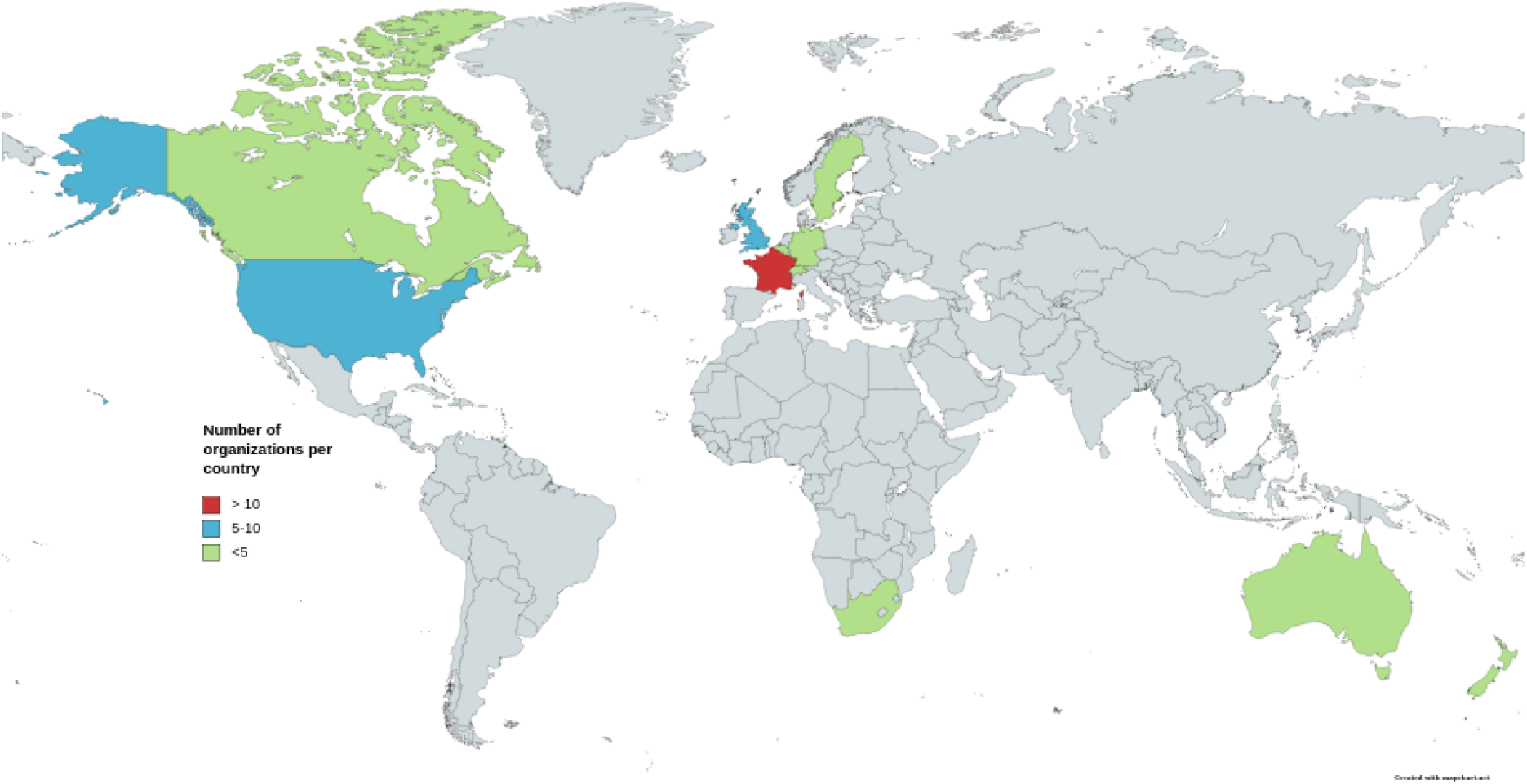
Number of organizations analyzed according to their countries

We now report the main observations that could be made during our analysis, topic by topic.

### Neutrality or reduction?

The reduction targets can be expressed in terms of carbon neutrality (with remaining emissions being offset), in terms of absolute or relative reduction, or both. We have observed two main cases (see Figure 2): GHG reduction targets with no mention of a neutrality objective (18 organizations), or reduction targets combined with a neutrality objective (26 organizations). Only 4 organizations did not give a precise target, and 3 had carbon neutrality targets without reduction targets.

**Figure 2.**
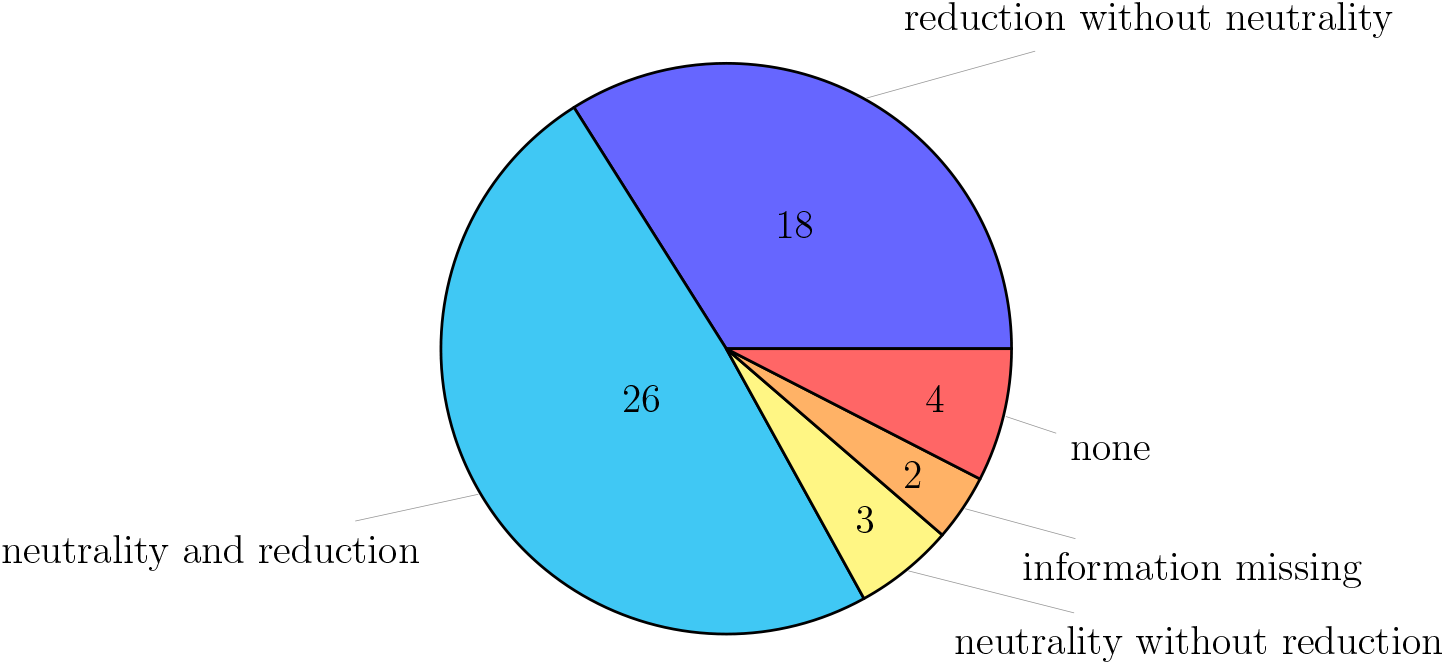
Number of organizations having adopted the following objectives among GHG reduction and neutrality

Achieving neutrality can be done by focusing on emission reduction or by increasing offsetting, but both solutions are not equivalent in terms of their effects on climate change; so organizations should ideally detail the emission reductions associated to a neutrality objective. Yet, few organizations clearly associate their neutrality objective with emission reductions, these two types of objectives often being for different timelines. And the role of offsetting, usually used to achieve neutrality, is not always clear, although 26 organizations use or plan to use offsetting in their strategy. Although the concept of “carbon neutrality” for an organization is debated, its limits were not discussed in the documents analyzed. We discuss this important issue in Section 3.

### Energy use

Most organizations take into account energy use (electricity and heat) in their GHG reporting, these emissions accounting for most of emissions of scopes 1 and 2. This corresponds respectively to 40 organizations both for electricity and heat, as can be seen from Figure 3. Since many organizations consider electricity and heat in a same “energy” emission source, we will not distinguish those in the rest of this paper.

**Figure 3.**
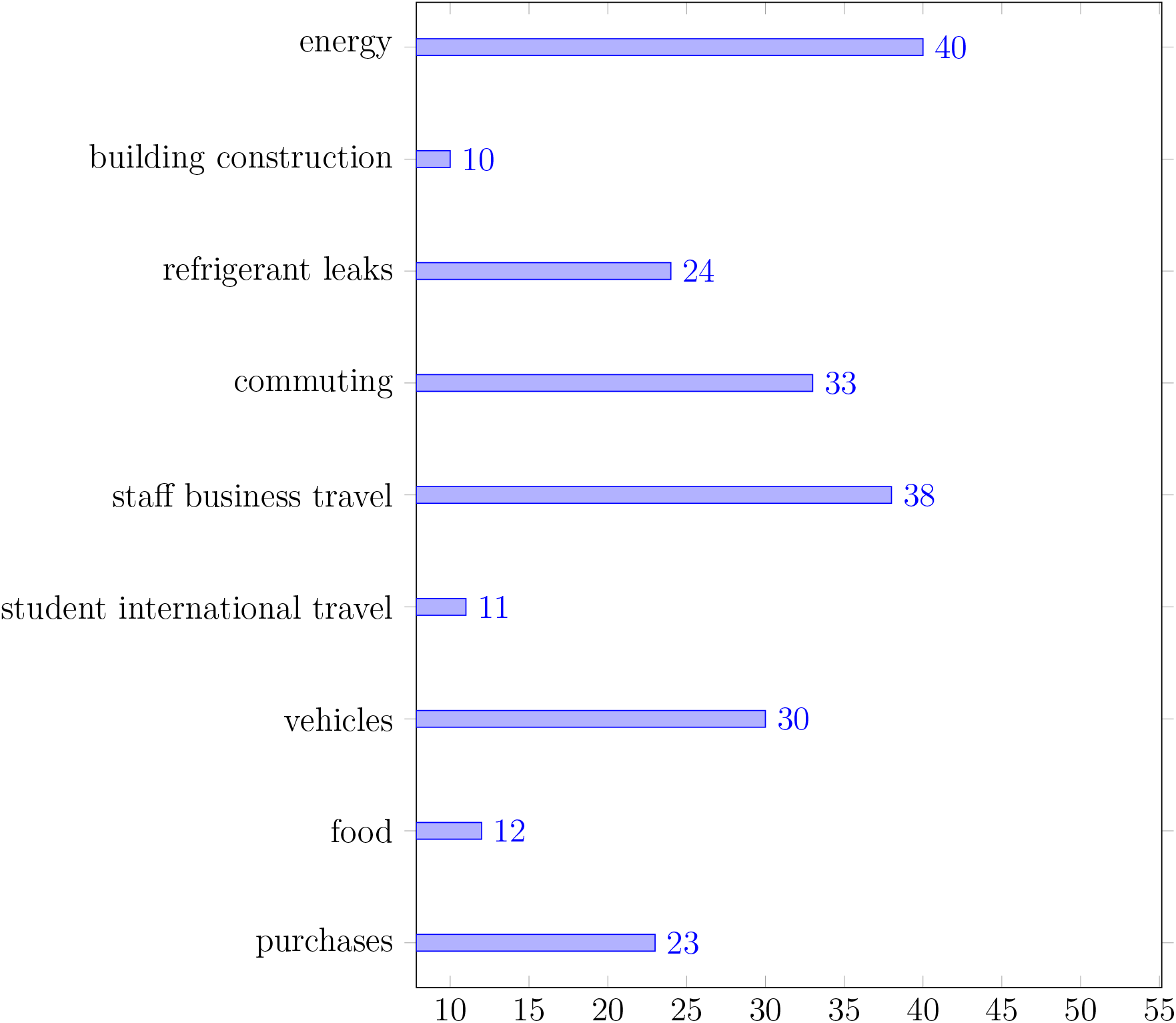
Emission sources taken into account in the existing carbon reports

We have noted though that most of the organizations considered do not specify the emission factor that they use to calculate their electricity-related carbon footprint, generally not even indicating whether it is a location-based or market-based factor. Only 10 organizations specify their emission factor, 5 using a location-based one and 5 using a market-based one; 6 more organizations also use a location-based emission factor that is included in the GES 1point5 tool used [Mariette et al., 2022]. The various choices made and the adopted methodology regarding this item matter and will be the subject of further discussion in Section 3.

Concerning the definition of targets, 20 organizations clearly express a reduction target for energy use, 12 in terms of GHG emissions, and 8 others in terms of energy consumption (for example the Ecole Centrale de Lyon aims at a 15% reduction of its energy consumption by 2025 and 40% by 2030) as can be seen in Figure 4. 10 organizations also define targets in terms of renewable energy share.

**Figure 4.**
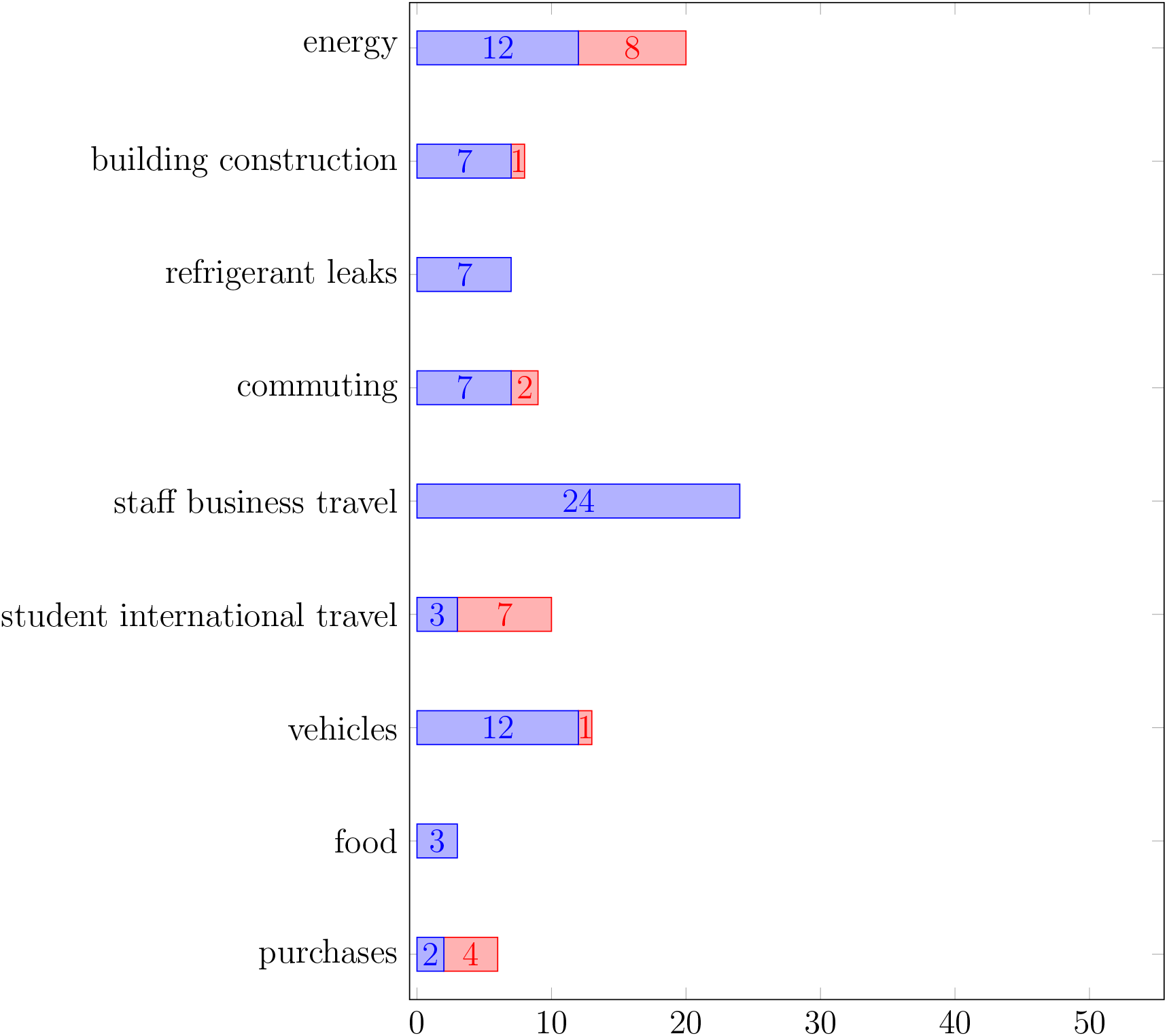
Emission sources for which explicit reduction targets are given: in blue the targets expressed in terms of GHG reduction, in red the targets expressed only in terms of activity data (for example energy consumption only)

### Building construction

Taking into account the construction of the buildings hosting the students and the employees of the organizations, as fixed assets during a given number of years, is considered by a minority of organizations: 10 according to Figure 3. Among these 10 organizations, 7 have defined target reductions expressed in GHG emissions to reduce the impact of building construction, while 1 other has expressed the target reduction in activity of building construction, as seen in Figure 4.

### Staff business travel

A majority of organizations, 38 from Figure 3, include business travel from their employees in their GHG emissions. In practice this includes teaching-related reasons - e.g., teaching abroad or signing of agreements - or research-related reasons - e.g., international panels, committees, conferences, project meetings, field studies… Usually the considered perimeter for business travel corresponds to budget lines financed by the organizations.

As the vast majority of emissions from these trips are due to air travel [Ben Ari et al., 2023], 6 organizations limit themselves to air travel only.

Few organizations specify the emission factors or the tool used to calculate the carbon footprint associated with such travel, with the exception of one organization which indicates using Myclimate ∥, and the French organizations using the GES 1point5 tool. This is problematic, as the emission factors associated with aviation factors can range from simple to double (or even triple), depending on whether or not condensation trails are taken into account [Lee et al., 2021].

Finally, reduction targets were found directly associated with this item or included in the scope 3 target. From Figure 4, it is seen that 24 organizations set a reduction target expressed in GHG emissions.

### Student international travel

In addition to staff business travel, undergraduate student study-related travel is a relevant emission item since international mobility (to and from the country) increased a lot in the past decades either through dedicated programs run by the organizations/countries or by individuals themselves (for example the number of participants in the Erasmus European mobility program has nearly doubled between 2014 and 2022 [Erasmus, 2023]). For this reason, not including travel by students largely underestimates the emissions from travel.

A minority of 11 organizations take into account this emission source in their GHG emissions (see Figure 3). Nevertheless we note that this is partly due to 19 organizations being research laboratories for which this item is not relevant. As for staff business travel above, the trips taken into account may be financed by the university or not. In particular, student travel not funded by the university will usually not be taken into account as it will not appear in university’s budget lines. Figure 4 shows that regarding this item a low number of 3 organizations set target reductions in their GHG emissions, while 7 organizations adopted quantitative activity reduction targets.

### Commuting

Commuting by students and staff is also a relevant GHG emission item. As for international student travel, not including commuting by students can largely underestimate the emissions from commuting. 33 organizations include commuting as shown in Figure 3, which is comparable to but smaller than the number of organizations considering staff business travel. Among these 33 organizations, Figure 4 shows that only 7 set GHG emissions reduction targets, while 2 set quantitative reduction targets but expressed in activity reduction of commuting.

### Purchases

Purchases is a broad, somewhat catch-all category, where organizations generally stash away what is not accounted for more precisely. For this reason, the perimeter of purchases can be very broad going for example from paper to furniture, numerical devices, consumables (both scientific and non-scientific ones), scientific instruments or also services (both scientific and non-scientific ones). The estimate of the carbon footprint of purchases is a highly delicate task. In the absence of a more precise method, it is usually done using monetary emission factors, which are less precise than physical emission factors (the latter being most often not available for purchases).

Not surprisingly, a minority of organizations indicate taking into account the carbon footprint of purchases in their GHG emissions reports. 23 organizations include purchases as shown in Figure 3. However, we note that the perimeters of purchases explicitly considered vary a lot between the organizations we have analyzed. For instance a significant number of organizations included only IT equipment. It may be noted that IT equipment emissions can be assessed using a physical approach. This is the case with the GES 1point5 tool, which is based on the Ecodiag tool *¶*. Others included separately paper, identified as a potential important item in GHG emissions. Among the French research organizations, many included the broad panel of purchases taken into account in GES 1point5 [De Paepe et al., 2023] which considers about 1,400 different categories of purchases. According to our analysis, only 2 organizations set GHG emissions reduction targets on purchases, while 4 others set reduction targets in terms of activity reduction (see Figure 4).

Several hypotheses could explain this very heterogeneous situation. First, including purchases in a GHG report is difficult due to the very large variety of goods and services that are involved. Relying on widely shared typologies of purchases, such as what was done in GES 1point5, is a way to cope with this difficulty. Second, the monetary methods used to estimate purchases emissions makes it difficult to implement their reduction in a scenario because of the high uncertainties of emission factors, and their lack of connection with actual environmental impacts: for example choosing a more sustainable alternative when buying a product may lead to an increased price, and, if the emission factor is not precise enough, to an increased estimation of GHG emissions. Third, reducing the emissions due to purchases will lead to very different solutions depending on the typology of purchases, so it is harder to come up with ready-made solutions that will work in all organizations.

The absence of purchases is a very important blind spot in GHG reporting and reduction plans of higher education and research organizations, as purchases in particular and scope 3 more generally dominate their emissions [De Paepe et al., 2023].

### Other emission sources

Other emissions sources were considered in the GHG emissions reports of the 53 studied organizations.

24 organizations included refrigerant leaks from cooling systems. This item is highly relevant for organizations having many air-conditioned rooms or labs, especially as the emission factors associated with these refrigerants are very high. In addition, 7 organizations set GHG emissions reduction targets on this item.

Food is an important source of GHG emissions. The perimeter of this item is rather broad including students and staff catering, but also delivered food or drinks ordered by organizations or students. However, as shown in Figure 3 a small minority of 12 organizations included food in their GHG reports. The mentioned perimeter is heterogeneous but very few organizations include explicitly students and staff catering. This can lead to a large underestimation of this item. For staff members, part of this item can be included in purchases but usually not for students. 2 organizations set GHG emissions reduction targets, while 4 set reduction targets in terms of activity. This emission source, like commuting, is at the interface between private and public spheres, but most probably has a large carbon footprint. Organizations have levers to reduce the corresponding GHG emissions. Thus, this item should be taken into account.

A majority of the studied organizations included the vehicles owned in their GHG reports: 30 as seen from Figure 3. Out of these, 12 set GHG emissions reduction targets, while 1 set reduction targets in terms of activity reduction.

It can be noted that no organization took into account (separately) their subsidies to large research infrastructures, participation of their staff to research projects involving large research infrastructures or their use of large computing facilities that may yet have a huge contribution to the carbon footprint of research [Knödlseder et al., 2022].

The items of water consumption and waste were considered by several organizations. The importance of these items does not express in terms of GHG emissions, which are low, but more in terms of general environmental footprint. For this reason, we did not include here these items although their importance is very high in the context of a general ecological crisis.

### Target years

As expected, many institutions have committed to reduction or neutrality targets for 2030: 31 organizations cite 2030 as a target year. Many institutions also have shorter-term intermediate targets, such as 2025 (16 organizations define targets for years between 2014 and 2026). This is in line with the Science Based Targets initiative, which recommends setting a short-term target (5 to 10 years) and a long-term target (2050 at the latest).

A few organizations have already achieved neutrality for Scopes 1 and 2, and commit to maintaining it.

Many organizations also made commitments for years beyond 2030, between 2040 and 2050, mostly regarding neutrality objectives.

### Reduction target

As mentioned before, it is very difficult to compare reduction targets which differ in terms of perimeter and deadlines.

By 2030, to take a date common to several organizations and a target year in climate commitments^+^, objectives vary widely from one organization to another: the University of Oregon in the USA, for example, has committed to a 34% reduction in emissions between 2019 and 2030, with emission sources in scopes 1, 2 and 3; the University of Tasmania in Australia has committed to a 50% reduction compared with 2015 emissions on numerous items (scopes 1 and 2 and several items in scope 3); the Swiss Federal Institute of Technology in Zurich, Switzerland, has committed to a 50% reduction in emissions by 2030 compared to 2006 levels (scopes 1, 2 and 3); the University of British Columbia in Canada has committed to an 80% reduction compared to 2007 levels on scopes 1 and 2 and 45% by 2030 compared to 2010 on scope 3. It should be remembered that scope 3 generally accounts for the vast majority of emissions, so even considerable reduction commitments based solely on scopes 1 and 2 are in reality very partial.

More generally, it can be noted that the reference years vary, probably depending on the first estimate of the organization’s GHG emissions. Nearly all organizations have set a reduction target based on a reference year more recent than 1990. The year 2019, the last pre-pandemic year, is often chosen as a reference. The year 2019 is also used in the IPCC’s 6th report (Mitigation section, [Shukla et al., 2022]) to express a reduction target consistent with keeping the global temperature increase below +1.5°C: -43% in 2030 compared with 2019, thus moving away from the 1990 reference.

The reduction percentages also vary, some being aligned with the low-carbon trajectories of the Paris Agreement or more demanding trajectories, but the reference years and items taken into account vary enormously, making it difficult to know whether these targets really fall within this framework.

### Trajectories

6 organizations only have defined complete reduction trajectories, 4 of them broken down by emission source, showing priorities in their climate policy as the reduction targets vary across emission sources. The other 2 focus on the expected gains due to some actions on total emissions, without showing the reduction variations by emission source. Four organizations adopt trajectories that reveal a scheduling of efforts with different rates of reduction between intermediate target years. The trajectories of the other two show a uniform reduction rate across time.

Only one organization has a constraint on cumulative emissions: Columbia University assesses that “Total cumulative emissions from 2019 to the date net zero is achieved shall not exceed 14.6 times the University’s 2019 emissions”.

## 3. Discussion and recommendations

As it has been observed for carbon footprint reporting [Robinson et al., 2015, Valls-Val and Bovea, 2021, ALLEA, 2022], climate targets of higher education and research organizations are very heterogeneous, which makes it difficult to assess their relevance.

Based on our observations and SBTi guidelines, we thus propose guidelines for higher education and research organizations that wish to set climate targets, hoping that it could help them and also help the harmonization between organizations.

### Type of target

When expressing a low-carbon target, we recommend to express objectives in terms of percentage reduction in GHG emissions, or in terms of absolute value of emissions, and not (only) in terms of carbon neutrality. We also recommend using the term “contribution to global neutrality” rather than “carbon neutrality”, as proposed by the Carbone 4 report [Dugast, 2020].

We also suggest presenting separately the reduction in GHG emissions due to the organization’s internal policy and offsetting, as also proposed in [Dugast, 2020], in order to highlight the emission reductions planned. As offsetting is subject to a great deal of criticism, particularly concerning its actual climate impacts^*^, it is recommended that organizations base themselves on emission reduction trajectories that do not require offsetting.

We also recommend clarifying whether a constraint has been placed on cumulative emissions between now and the target year, and not just on the emissions target.

### Perimeter

Concerning the perimeter, we recommend to specify the scope as precisely as possible: provide as precise a list as possible of both the scopes and emission sources included in the reduction target, and those not included. For targets concerning business travel, clearly specify the scope for both staff and students: travel financed by the establishment only, incoming and outgoing international mobility, etc.

We also recommend ensuring that all potentially significant emission sources have been taken into account, and trying to estimate the proportion of the perimeter excluded. The most emitting sources are generally the following: purchases (including IT equipment), energy consumption (electricity and heating), business travel and commuting by students and staff, food, and building construction. Organizations with major cooling systems (datacenters, air-conditioned experimental laboratories, etc.), should take refrigerant leaks into account. Organizations with other specific features should take into account these specific sources, for example subsidies for research platforms or large instruments, or computation time on shared infrastructures.

The climate targets are usually based on a GHG report, but this report is not always explicitly referred to in the documents: we recommend including the GHG report or a link to the GHG report before defining the targets.

### Methodology

Organizations should specify the emission factors, methodology or tools used: for all sources, specify whether the emission factor is a physical or monetary one; for targets concerning energy consumption in particular (scope 2), specify the source and type of emission factor used (in particular market-based or location-based); for the emissions due to air travel, specify the emission factors used and whether or not condensation trails are taken into account. We suggest to take condensation trails into account when estimating the impact of air travel, since non-CO_2_ impact of air travel is high.

### Deadlines

Following the SBTi guidelines, we suggest committing to close and later deadlines. As 2030 is a target date set by the Paris Agreement, it is advisable to refer to it. As the impact of reductions is not the same depending on the trajectory, since the accumulation of emissions is also crucial, it is useful to also set intermediate targets.

The question of the long-term trajectory, in 2040 or 2050 (as in the Paris Agreement), remains useful for anchoring the organization’s transition objective.

### Reduction targets

Even though the reference year may be chosen due to practical reasons (first GHG reporting for example) more than climate-related ones, the organizations should specify and justify the reference year used. If possible, they should chose a year that can be linked to IPCC scenarios, such as 2019. This will enable them to indicate whether the reduction targets are in line with the IPCC scenario of warming below 1.5°, whose reduction target for 2030 compared with 2019 is 43% [Shukla et al., 2022], and explain the reasons for a lower or more ambitious target.

It is also important to define targets in terms of GHG emissions and not only activity data (energy use in kWh for example): the emission factors may also evolve, which will influence the GHG emissions.

### Trajectory

We recommend developing strategic thinking and trajectories to allocate the reduction effort to each of the emission sources, spread the reduction effort over time and set intermediate targets.

A trajectory should reflect the organization’s strategic choices, by enabling intermediate targets to be visualized for each emission source. As the strategy must determine when measures are to be implemented, the curves will probably not be linear. Trajectories guide both the construction of an action plan and the monitoring of its execution. They are also essential to compute cumulative emissions and show how a target in terms of carbon budget can be met, if such an ambitious objective has been set.

Finally, organizations need to regularly revise and update their reduction targets and trajectories, based on regular GHG reporting.

## 4. Conclusion

In this paper, we address the question of how climate targets could be set by higher education and research organizations. We present a review of documents from more than 50 organizations worldwide that set climate targets. More and more universities and laboratories publish their GHG reports and actions taken to reduce their impacts. Nevertheless, our analysis shows that they present a very large heterogeneity in terms of covered perimeter (i.e. scopes and categories), deadlines, type of objectives (i.e. carbon neutrality, net zero, or GHG emissions reduction), reduction percentage, role of offsetting, absence or presence of projected trajectories.

There is an urgent need to provide good practice protocol so that all the organizations can build and set their own objectives using a common and comprehensive framework of definitions and actions. In order to help higher education and research organizations define their climate targets and trajectories, we propose explicit guidelines to deal with these elements. Our recommendations highlight the need to specify, explain and detail all the choices and parameters used at every stage, and to adapt the general cross-domain guidelines of the Science Based Targets initiative to the specifics of this sector. The approach developed here is quite generic and could also be useful for other public institutions aiming at setting climate targets and trajectories.

## 5. Methods

### 5.1. Identification of the documents

In a first step, we analyzed existing climate strategies of higher education and research organizations. The organizational perimeters considered are of different natures: although most documents concern universities, we also considered research laboratories or institutes, for example. For simplicity’s sake, we will refer to “organizations” in all cases hereafter.

To carry out this analysis, we searched for sustainable development policy documents in English or French. Two types of documents generally exist: documents presenting the organization’s general policy on this subject, which are public; and specific documents including action plans, which are not always public. We relied mainly on public documents, except for certain laboratories participating in the Labos 1point5 collective *♯*, for which we had more information at our disposal via the Labos 1point5 experimentation scheme.

In order to identify relevant documents, we proceeded in two steps:

- We started with organizations that we already knew about, because we work there, have colleagues there, or had already read about their low-carbon commitments.
- We extended this list in a number of ways: searching for academic institutions by geographical area, and studying institutions that ranked highly in relevant rankings such as the Times Higher Education Impact Rankings.

We selected documents if they included a detailed GHG reduction strategy only, and ended up with documents for 53 organizations.

As this study was done in 2022, the documents analyzed are those available at that time, and some organizations may have made progress since then.

### 5.2. Document analysis methodology

In order to define GHG reduction targets, organizations should at least include:

- An organizational perimeter.
- One or several target years.
- One or several target reductions (percentage of reduction or neutrality).

The perimeter should define the scopes and GHG emission sources considered for the reduction target, generally based on GHG reporting. Two main standards are used for this: the GHG Protocol and the ISO 14069 norm. Table 1 summarizes the main GHG emission sources for higher education and research organizations and the associated scopes and categories in the GHG Protocol and ISO 14069 norm. For each emission source, further information may be required: for example, for commuting, are both students and staff considered? Table 2 provides details on the information that was searched for in the documents.

**Table 1.** Main GHG emission sources for higher education and research organizations, and correspondence with GHG Protocol and ISO 14069 scopes and categories.

| Emission sources | GHG protocol | ISO 14069 |
| --- | --- | --- |
| fuel for the organization's vehicles | 1 | 1 |
| electricity | 2 (or 1) | 2 (or 1) |
| heating | 2 (or 1) | 2 (or 1) |
| cooling (refrigerant leaks) | 2 | 2 |
| commuting (students & staff) | 3 | 3 |
| business travel (students & staff) | 3 | 3 |
| purchases of products and services | 3 | 4 |
| capital assets, in particular building construction |  |  |

**Table 2.** Information searched for in the documents analyzed, by emission source.

| Emission source | Further information |
| --- | --- |
| electricity | emission factor given?<br>location-based or market-based emission factor? |
| business travel | students and/or staff?<br>flights only?<br>financed by the organization only?<br>emission factors given? |
| commuting | students and/or staff? |

Concerning the target year, we expected to find commitments for the timeframes corresponding to the Paris Agreement, i.e. 2030, which is moreover quite close, and 2050. These target years would also be in line with the Science Based Targets initiative that calls for a short-term target (5 to 10 years) and a long-term target (2050 at the latest).

Concerning the target reductions, it is obviously difficult to consider percentages of reduction envisaged independently of the initial carbon footprint of the organizations and their specific characteristics (thematic, geographical, budget,…). Furthermore, it is very difficult to compare percentages that are strongly linked to different reference years, deadlines and perimeters used. It is nonetheless interesting to consider their homogeneity and compatibility with the Paris Agreement. The SBTi cross-sectors absolute reduction approach prescribes a 4.2% (resp. 2.5) minimum linear annual rate of reduction for base year of 2020 or earlier *††* and scopes 1 and 2 (resp. 3), leading to a 42% (resp. 25) minimum reduction in emissions by 2030, from 2020 levels.

Commitments to low-carbon objectives can be expressed in terms of GHG emission reduction, or in terms of “carbon neutrality”, “climate neutrality” or “net zero [emissions]”. However, expressing targets in terms of reduction or neutrality is not equivalent: A carbon-neutrality objective can be achieved either by reducing emissions or by increasing offsetting. The concept of carbon neutrality for organizations has the following limitations, as detailed in [Dugast, 2020]: possible variations in scopes, invisibility of the emission reduction, impossibility to apply it universally… [Dugast, 2020] recommends talking about a “contribution to global neutrality” instead of a “carbon neutrality” target. The SBTi guidelines enforce the specification of many of these points, as explained above, leading to more relevant “net zero” targets.

In order to correspond to actual possible reductions, the overall target should be broken down by emission source and over time, and reduction trajectories determined. These trajectories are the numerical expression of strategic choices: on which emission sources do the organizations wish to take the strongest actions, within what timeframe, and with which intermediate targets. For most of the emission sources, the target reductions can be expressed either in terms of GHG emissions or in terms of activity data. For example, a reduction target for the emissions due to electricity use may be expressed in terms of GHG emissions, or in terms of energy consumption (in kWh for example). This is not equivalent, since the emission factor, that enables to convert the activity data into GHG emissions, may also change during the period considered. In our document, we will focus on targets expressed in terms of GHG emission reduction, but will also mention other possibilities.

Trajectories are important since they determine whether the organization remains under a given carbon budget, i.e. cumulative emissions that must not be exceeded. With a constant final target, the faster the reduction is in the first few years, the more the carbon budget is preserved.

Strategic consideration of the distribution of effort by emission source and over time is a political matter, but also depends on the existence of levers for action. For example, a laboratory may have more levers for action on business travel and purchasing emissions than on building-related emissions, since this emission source is more likely to depend on the establishments directly and not on the laboratories.

In a scenario-based method, the organization must first define a trajectory, in order to characterize the organization’s low-carbon policy and guide the choice of measures. This initial trajectory may be adjusted when reduction actions with estimated reduction effect are defined.

In a scenario-based method, it is also important to try and imagine several narratives characterizing the organization’s envisaged low-carbon policies and associated trajectories in order to make an informed strategic choice, but as no narrative was present in the documents we studied, we focus here on the quantitative objectives of the adopted trajectory only.

## Acknowledgments

Portions of this article were written by post-editing text that was translated from French into English with www.DeepL.com/Translator (free version).

The authors thank Audrey Sabbagh and Olivier Ragueneau for initiating this work. We also thank Jürgen Knödlseder and Malo Serra for collecting documents and participating to a few meetings, and Marion Avet, Andrá Estevez-Torres, Ivan Magrin-Chagnolleau, Matthieu Romagny and Cáline Serrano for their feedbacks on the first version of the paper.

## Supplementary material

The documents that we analyzed are available in the following directory: https://cloud.le-pic.org/s/mLicnzZoCrWt95r

## Author contributions

All authors contributed to the data collection from the organizations documents. L.P. wrote scripts to automate the counts. A.-L.L. wrote the first draft of the paper, C.B. made substantial refinements. A.-S.M., L.P., E.J., A.M., B.D. and L.V. made edits and critical revisions. All authors approved the final manuscript.

## Footnotes

‡ https://sciencebasedtargets.org/

§ https://labos1point5.org/

∥ https://www.myclimate.org/en/

¶ https://ecoinfo.cnrs.fr/ecodiag-calcul

+ such as the Paris Agreement

* https://www.theguardian.com/environment/2023/jan/18/revealed-forest-carbonoffsets-biggest-provider-worthless-verra-aoe or https://www.science.org/doi/10.1126/science.adj6951 for example

♯ https://labos1point5.org/

†† which was mainly the case due to the dates of our corpus

## Notes

https://cloud.le-pic.org/s/mLicnzZoCrWt95r

### Competing Interest Statement

The authors have declared no competing interest.

